# Cystatin C loaded in brain-derived extracellular vesicles rescues synapses after ischemic insult *in vitro* and *in vivo*

**DOI:** 10.1101/2023.06.16.545303

**Authors:** Yuqi Gui, Yohan Kim, Santra Brenna, Maximilian Wilmes, Giorgio Zaghen, Lennart Kuchenbecker-Pöls, Hannah Voß, Antonia Gocke, Hartmut Schlüter, Michaela Schweizer, Hermann C. Altmeppen, Tim Magnus, Efrat Levy, Berta Puig

## Abstract

Synaptic loss is an early event in the undersupplied but not yet irreversibly injured penumbra area after an ischemic stroke. Promoting synaptic preservation in this area would likely improve functional neurological recovery. In the present study, we aimed to detect proteins involved in endogenous protection mechanisms of synapses in the penumbra after stroke and to analyse the potential beneficial effect of these candidates for a prospective stroke treatment. For this, we performed Liquid Chromatography coupled to Mass Spectrometry (LC-MS)-based proteomics of synaptosomes isolated from the ipsilateral hemispheres of mice subjected to experimental stroke at different time points (24 h, 4 and 7 days) and compared them to sham-operated mice. Proteomic analyses indicated that among the differentially expressed proteins between the two groups, cystatin C (CysC) was significantly increased at 24 h and 4 days following stroke, before returning to steady-state levels at 7 days, thus indicating a potential transient and intrinsic rescue mechanism attempt of neurons. When CysC was applied to primary neuronal cultures subjected to an *in vitro* model of ischemic damage, this treatment significantly improved the preservation of synaptic structures. Notably, similar effects were observed when CysC was loaded into brain-derived extracellular vesicles (BDEVs). Finally, when CysC contained in BDEVs was administered intracerebroventricularly to stroked mice, it significantly increased the expression of synaptic markers such as SNAP25, Homer-1, and NCAM in the penumbra area compared to the group supplied with empty BDEVs. Thus, we show that CysC-loaded BDEVs promote synaptic protection after ischemic damage *in vitro* and *in vivo*, opening the possibility of a therapeutic use in stroke patients.

## INTRODUCTION

Ischemic stroke is a condition characterized by high mortality and a high disability rate [1]. It is caused when an embolus or thrombus occludes a cerebral vessel leading to immediate cerebral tissue damage. This is followed by a complex pathophysiological reaction involving the activation of different cell types and neuroinflammation progressing in a spatiotemporal manner [2]. The core of the infarcted tissue (where cells rapidly die by oncosis/necroptosis) is surrounded by a hypoperfused area (the penumbra), where neurons are functionally damaged but still rescuable [3]. Currently approved therapies, such as the delivery of recombinant tissue plasminogen activator (rtPA) or mechanical thrombectomy (in case of a large arterial occlusion), are aimed at reperfusing the affected area as soon as possible to rescue cells in the penumbra. Both therapies need to be performed within a critical period, originally between 4.5 to 6 h after stroke onset, although new studies show that the time window can be extended if certain inclusion criteria are met [4–6]. Beyond this critical time window, these treatments are even detrimental as they increase the risk of hemorrhagic transformation [7]. As a consequence of this limited treatment window, strict inclusion criteria, and a general lack of proper devices and expertise for thrombectomy procedures out of big hospitals, the majority of stroke patients cannot benefit from these interventions [8–10]. Therefore, identifying (endogenous) protective mechanisms and exploring new regenerative therapies to promote neuronal recovery in the penumbra that are independent of the above-mentioned limitations is of utmost importance.

Synaptic neurotransmission is highly dependent on energy supply, and the lack of the latter is one of the first consequences of vessel occlusion and hypoxia. Both, pre- and post-synaptic terminals are rapidly lost after ischemia. In the penumbra, this process may occur without neuronal loss as neurons enter an electrical silence to save energy but are still viable and metabolically active [11]. *In vitro* studies show that synaptic connectivity can be fully restored 6 h after hypoxia, implying the reestablishment of neuronal functionality [12, 13]. Moreover, synaptic reconstruction is a feature seen in the penumbra during the early phases of recovery after stroke. Thus, in a gerbil model of stroke, the number of synapses initially decreased up to day 4, then started recovering one week after stroke [14], and increased SNAP-25 (a synaptic marker protein involved in axonal outgrowth) immunoreactivity was observed as early as two days after ischemic induction [15]. *In vivo* experiments revealed that, after moderate ischemic damage (equivalent to damage in the penumbra area), dendritic spines were resilient for the first 5 h, to be gradually lost after 7 h, triggering the activation of signalling cascades that promoted apoptotic cell death [16].

Rescuing synapses at the penumbra, increasing connectivity and functionality (and thereby neuronal survival), may represent an important therapeutical niche [17, 18]. In the present study, we aimed to unveil protein changes undergone by synapses at different time points after stroke to identify candidates that may be involved in the intrinsic mechanisms of synaptic protection and/or recovery. Our findings show that cystatin C (CysC), a cysteine protease inhibitor expressed by all nucleated cells, is increased in synaptosomes isolated from mice at early time points after stroke. We demonstrate that CysC rescues synapses *in vitro* when externally delivered either free or loaded into brain-derived extracellular vesicles (BDEVs). Furthermore, treatment of stroked mice with CysC-loaded BDEVs increased the expression of synaptic marker proteins as early as 24 h after the insult. Thus, due to the capability of EVs to cross the blood-brain barrier (BBB) and the higher stability of drugs when encapsulated by EVs [19], we envisage CysC-loaded EVs as a potential clinically relevant future treatment for stroke.

## MATERIALS AND METHODS

### Animals

The mice used for the stroke experiments were male C57BL/6 aged between 11 and 17 weeks, kept under a 12 h dark-light cycle with *ad libitum* access to food and water. Animal experiments were approved by the local animal care committee (*Behörde für Gesundheit und Verbraucherschutz, Veterinärwesen und Lebensmittelsicherheit* of the *Freie und Hansestadt Hamburg*, project number N045/2018 and ORG1005) and in compliance with the guidelines of the animal facility of the University Medical Center Hamburg-Eppendorf. BDEVs were isolated from the brain of 4 months old CysC knock-out mice [20]. Animal procedures were performed following the National Institutes of Health guidelines with approval from the Institutional Animal Care and Use Committee at the Nathan S. Kline Institute for Psychiatric Research.

### Transient middle cerebral artery occlusion (tMCAO)

Left-side tMCAO in mice was performed as previously described [21] with minor changes. Briefly, after anesthesia and analgesia, a one-centimeter incision was made along the mouse neck midline, and the left carotid artery branch was exposed. The external carotid artery was ligated and notched at the distal end, and a filament (6-0 nylon, 602312PK10, Doccol) was inserted to block the middle cerebral artery. After 45 min of arterial occlusion, the filament was removed to allow reperfusion. The sham-operated mice also had the arteries exposed but no occlusion was made.

### Isolation of synaptosomes from mouse brains

Synaptosomes were isolated from fresh brain tissue and an overview of the whole procedure is depicted in Fig. 1A. Briefly, the ipsilateral hemisphere was excised and gently homogenized with a Dounce grinder by 15 strokes in the homogenization buffer (HB: 0.32M sucrose in 5mM HEPES pH = 7.4 supplemented with protease inhibitors (P.I., Sigma #11697498001) at a ratio of 1:10 w/v on ice. Samples were then centrifuged at 1,000×*g* for 10 min at 4°C; the supernatant (S1) was kept on ice, and the pellet (P1) was again resuspended in HB buffer (1:10 w/v). The resulting supernatant (S2) was mixed with S1 and centrifuged at 12,000×*g* for 15 min at 4°C. The resulting pellet (P3) was resuspended again in the HB buffer (1:10 w/v) and centrifuged at 12,000×*g* for 20 min at 4°C. The resulting pellet (P4, crude synaptosome fraction) was resuspended in 1 mL of isolation buffer (IB: 0.2M sucrose in 5mM HEPES buffer pH = 7.4 supplemented with P.I.) and layered on top of a sucrose gradient (0.8M, 1.0M, 1.2M sucrose in 5mM HEPES buffer pH = 7.4). Samples were then centrifuged at 82,600×*g* at 4°C for 2 h (Beckman ultracentrifuge optima L-100 XP). After the first ultracentrifugation, the white band between the 1.0M and 1.2M sucrose layers was collected, resuspended in 10 mL of IB, and centrifuged again at 26,000×*g* at 4°C for 30 min. The final synaptosome pellet was resuspended either in 4% paraformaldehyde (PFA) containing 2.5% glutaraldehyde for electron microscopy or in 10mM HEPES buffer pH = 7.4 supplemented with P.I. for mass spectrometry or western blot analysis. For comparison purposes, 100 µL of total brain homogenate were extracted and mixed with 100 µL of 2× radio-immunoprecipitation assay (RIPA) buffer (50mM Tris-HCl pH = 7.4, 150mM NaCl, 1% NP40, 0.5% Na-Deoxycholate and 0.1% SDS).

**Figure 1.**
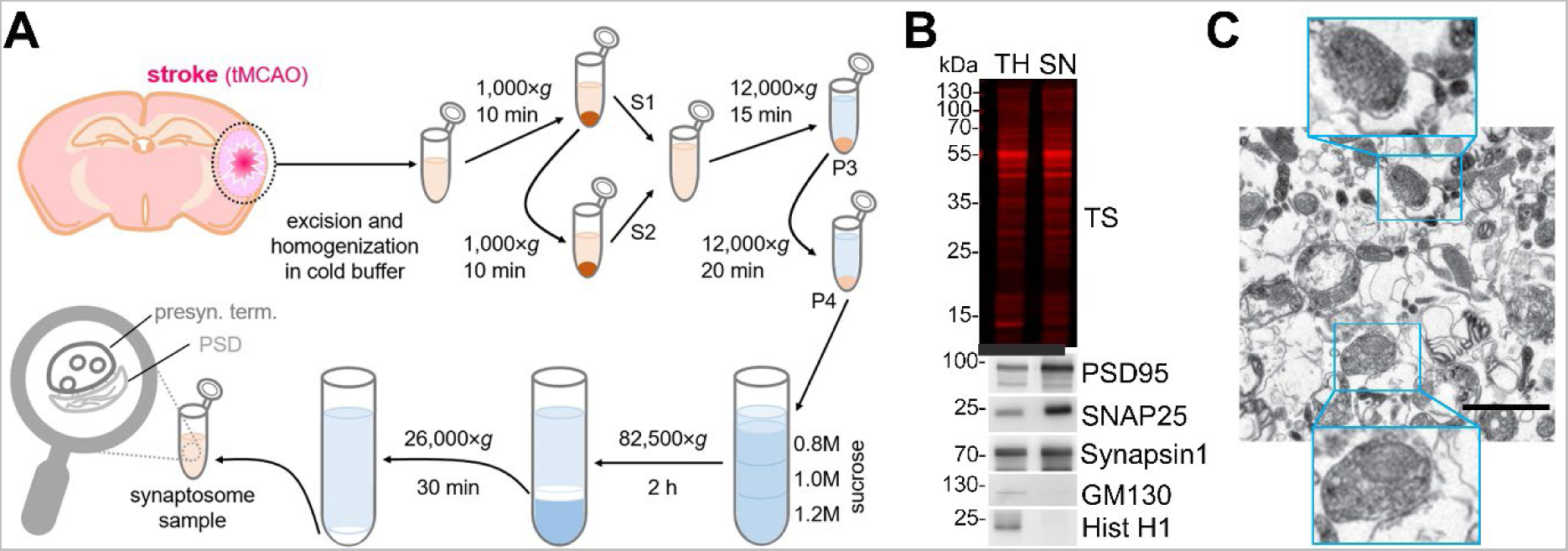
Synaptosome isolation and characterization. (A) Procedure overview. The stroke core and the penumbra tissue from mouse brains were dissected and homogenized in cold homogenization buffer, and subjected to a series of differential centrifugation and sucrose gradient ultracentrifugation to finally yield purified synaptosome samples. (B) Western blot analysis of the synaptosomal (SN) fraction. Total protein staining (TS) shows that equal amounts of proteins were loaded. TH is a total brain homogenate loaded for comparison purposes. Enrichment of synaptic proteins (PSD95, SNAP25 and Synapsin1) and absence of contaminations (GM130 as a Golgi marker protein and HistH1 as a nuclear marker) in the SN fraction shows the performance of the isolation procedure. (C) Representative electron microscopy picture of the final preparation showing several synaptosome structures (highlighted in the blue squares). The scale bar is 1 µm.

### Electron microscopy

Synaptosome pellets were fixed in 4% PFA containing 2.5% glutaraldehyde, centrifuged, washed three times with PBS, embedded in 3% agarose, and cut into small slices. Osmification was performed on the slices to add more contrast to the samples. Afterwards, samples were gradually dehydrated with ethanol and embedded in Epon resin (Carl Roth, #2545.1) for sectioning, and trimmed into ultrathin sections (60 nm) with a Leica Ultra Cut. Samples were analysed using an EM902 transmission electron microscope (Zeiss, Germany) equipped with a 2K digital camera with a CCD lens and running ImageSP software (Tröndle).

### Electrophoresis and western blotting

Protein concentrations were measured with Pierce BCA protein assay (Thermo Scientific #23225) according to the manufactureŕs protocol. Samples were diluted to 1 µg/µL in Laemmli loading buffer 1× (60mM Tris-HCl, 2% SDS, 10% glycerol, 0.05% bromophenol blue, 1.25% β-mercaptoethanol, pH = 6.8) and deionized H_2_O, denatured at 95°C for 5 min, and 9 to 10 µg of protein were loaded in a 12% acrylamide gel. Electrophoresis was performed at a constant current of 150 mV in a Mini-PROTEAN Tetra cell (BIO-RAD). Proteins were then transferred onto nitrocellulose membrane (LI-COR Biosciences) in a wet chamber (Mini-PROTEAN II cell (BIO-RAD)) filled with transfer buffer (25mM Tris base, 192mM glycine, 10% methanol). The membranes were then stained with Revert Total Protein Stain Kit (LI-COR Biosciences) following the manufacturer’s instructions and images of the total staining were taken in an Odyssey® CLx Infrared Imaging System (LI-COR Biosciences). Subsequently, unspecific binding was blocked with Roti-Block buffer (Carl Roth, #A151.2) 1× for 1 h at room temperature (RT). Primary antibodies were incubated at 4°C overnight or over the weekend on a shaking platform. The antibodies used here were: mouse against PSD95 (1:1,000; Millipore #MABN68), mouse against GM130 (1:1,000; BD Biosciences #610822), rabbit against Synapsin1 (1:1,000; Synaptic Systems #163103), rabbit against SNAP25 (1:2,000; Cell Signaling #3926), mouse against Histone H1 (1:500; Millipore #MABE446), and goat against CysC (1:1,000; R&D #AF1238). Afterwards, the membranes were washed with 1× TBST buffer (10mM Tris Base, 14mM NaCl, pH = 7.4, 0.01% Tween-20) for 5 min three times and then incubated with the appropriate secondary antibody (anti-mouse #7076 or anti-rabbit #7074 (both 1:1,000; Cell Signaling) or anti-goat (1:1,000; Promega #V805A)) for 1 h at RT while shaking. The chemiluminescence signal was developed with SuperSignal West Pico PLUS chemiluminescent substrate (Thermo Scientific #34577) or SuperSignal West Femto maximum sensitivity substrate (Thermo Scientific #34095) and visualized with either a ChemiDoc imaging system (BIO-RAD) or an Azure 400 visible fluorescent western system (Biozym). The protein bands’ signal intensities were analysed with Image Studio software (LI-COR).

### Bottom-up proteomic LC-MS/MS measurement of murine synaptosomes

To detect proteomic differences of synaptosomes between the stroked and sham-treated mice, the protein concentrations of synaptosome samples resuspended in HEPES buffer were tested by the Pierce BCA Protein Assay Kit (Thermo Fisher #23225) following the manufacturer’s instructions and then diluted to 1 µg/µL. 20 µg of each sample were used for tryptic digestion. Disulfide bonds were reduced, using 10mM DTT for 30 min at 60°C. Alkylation was achieved with 20mM iodoacetamide (IAA) for 30 min at 37°C in the dark. Tryptic digestion was performed at a trypsin: protein ratio of 1:100 overnight at 37°C and stopped by adding 1% formic acid (FA). The sample was dried in a vacuum concentrator (SpeedVac SC110 Savant;Thermo Fisher Scientific) and stored at −80°C until further usage.

Directly before LC-MS analysis, samples were resolved in 0.1% FA to a final concentration of 1 µg/µL. 1 µg of tryptic peptides was injected into a UPLC (nano Ultra-Performance Liquid Chromatography, nanoAcquity system; Waters).

Chromatographic separation of peptides was achieved with a two-buffer system (buffer A: 0.1% FA in water, buffer B: 0.1% FA in acetonitrile (ANC)). Attached to the UPLC was a peptide trap (180 μm × 20 mm, 100 Å pore size, 5 μm particle size, Symme try C18; Waters) for online desalting and purification followed by a 25 cm C18 reversed-phase column (75 μm × 200 mm, 130 Å pore size, 1.7 μm particle size, Peptide BEH C18; Waters). Peptides were separated using an 80 min gradient with linearly increasing ACN concentration from 2% to 30% ACN in 65 min. The eluting peptides were analyzed on a Quadrupole Orbitrap hybrid mass spectrometer (QExactive; Thermo Fisher Scientific). Here, the ions being responsible for the 12 highest signal intensities per precursor scan (1 × 106 ions, 70,000 resolution, 240 ms fill time) were analyzed by MS/MS higher-energy collisional dissociation (HCD) at 25 normalized collision energy (1 × 105 ions, 17,500 Resolution, 50 ms fill time) in a range of 400-1,200 *m*/*z*. A dynamic precursor exclusion of 20 s was used.

### Raw data processing for proteome data

LC-MS/MS data were searched with the CHIMERYS algorithm integrated into the Proteome Discoverer software (v 3.0; Thermo Fisher Scientific), using inferys 2.1 fragmentation as a prediction model, against a reviewed murine Swissprot database, obtained in February 2023, containing 20,365 entries. Carbamidomethylation was set as a fixed modification for cysteine residues and the oxidation of methionine was allowed as a variable modification. A maximum number of 2 missing tryptic cleavages was set. Peptides between 7 and 30 amino acids were considered. The fragment tolerance was set to 20 ppm. A strict cutoff (false discovery rate (FDR) < 0.01) was set for peptide and protein identification. For target decoy selection, the Percolator node was used. Protein quantification was carried out using the Minora Algorithm implemented in Proteome Discoverer. For quantification, razor and unique peptides were used. Protein quantification was performed based on summed abundances. Data normalization was performed at the peptide level to the total peptide amount. Peptide and protein abundances were scaled to an average of 100. For data processing and statistical analysis, each time point (24 h, 4 days, 7 days) was handled separately.

### Primary neuronal cell culture

Hippocampal neurons were obtained from postnatal day 1 or 2 (P1-P2) mouse pups. After decapitation, the brains were extracted and cleaned from the meninges with caution. The hippocampi were collected and transferred to ice-cold 10mM glucose in PBS, which was later replaced with 5 mL digestion solution (PBS/10mM glucose, 0.5 mg/mL papain (Sigma #P4762-100MG) and 20 µg/mL DNase (Sigma #DN25- 10MG)). The digestion was performed for 30 min in a 37°C shaking water bath. The hippocampi were washed 4 times with plating medium (70% (v/v) MEM (Gibco #51200-046), PBS/20mM glucose, and 10% (v/v) horse serum (Gibco #16050-130)), gently dissociated with a pipette and resuspended in 1 mL of plating medium. Cell counting was conducted by adding 20 µL of the cell suspension to 20 µL Trypan Blue (Sigma #T8154-100ML), and cell numbers were determined with the Countess® II cytometer (Thermo Fisher Scientific). 70,000 cells/well were plated onto 13 mm diameter coverslips in 24-well plates containing plating medium (500 µL/well). The coverslips were previously coated with poly-L-lysine (Sigma #P9155-5MG) at 37°C for 2-3 h and washed three times with PBS. 3-4 h after plating, the plating medium was changed to maintenance medium (Neurobasal A (Gibco #10888-022) containing 0.1% Gentamicin (Thermo Fisher #15750-060), 1× Glutamax (Thermo Fisher #35050-038) and 2% B27 serum (Gibco #17504-044)) (1 mL/well). 24 h after plating, 20mM 5-Fluoro-2’-deoxyuridine (EMD Millipore #34333) was added to each well to eliminate non-neuronal cells. Half of the maintenance medium was replaced every 3-4 days with fresh media. Primary neurons were cultured for 14 days to allow them to establish a mature synaptic network.

### Isolation of BDEVs and CysC loading

Isolation of BDEVs was performed as published previously [22]. Briefly, frozen right hemibrains of mice were minced and incubated with 20 units/mL papain (Worthington) in Hibernate A solution (HA, 3.5 ml/sample; BrainBits) for 15 min at 37°C for gentle dissociation. The papain was inactivated by the addition of 6.5 mL of ice-cold HA supplemented with protease inhibitors [5 μg/mL leupeptin, 5 μg/mL antipain dihydrochloride, 5 μg/mL pepstatin A, 1mM phenylmethanesulfonyl fluoride (PMSF), 1μM E- 64, all from Sigma-Aldrich]. The digested brain tissue was centrifuged at 300×*g* for 10 min at 4°C to discard undigested tissue and intact cells. The supernatant was sequentially filtered using a 40 μm mesh filter (BD Biosciences) and a 0.2 μm syringe filter (Corning Life Sciences). The filtrate was subjected to sequential centrifugations at 4°C, at 2,000×*g* for 10 min and 10,000×*g* for 30 min to discard membranes and debris, and at 100,000×*g* for 70 min to pellet the EVs. The pellet was washed once in PBS, recentrifuged at 100,000×*g* for 70 min at 4°C and resuspended in 1.5 mL of a 40% (v/v) OptiPrep (iodixanol) solution, containing 10mM Tris-HCl (pH = 7.4), 0.25M sucrose, and 40% iodixanol (all reagents from Sigma-Aldrich). An OptiPrep density step gradient column was set up by placing the 40% OptiPrep-equilibrated BDEVs in the bottom of a column tube and carefully layering on the top of it a decreasing scale of OptiPrep solutions (1.5 mL steps for 20, 15, 13, 11, 9, and 7%, and 2 mL for 5%). The sucrose step gradient was centrifuged at 200,000×*g* for 16 h at 4°C. Fractions 3 to 6 were combined and centrifuged at 100,000×*g* for 70 min, followed by resuspension in PBS.

CysC protein was loaded into BDEVs by sonication [23]. The BDEV suspension (combined fractions 3-6) was quantified using a nanoparticle analysis machine and diluted to 4.8 × 10^9^ particles/µL. Human urine-derived CysC (Millipore Sigma #240896-50UG) was reconstituted in 500 µL of 100mM sodium acetate buffer, pH = 4.5, to reach a concentration of 0.1 µg/µL. The 100 µL BDEV suspension, which corresponds to approximately 100 µg of total BDEV protein, was mixed with 20 µL of CysC solution (2 µg) to yield a 50:1 ratio of BDEV protein to CysC in a total 1 mL of PBS. The mixture of BDEVs and CysC rotated overnight at 4°C. Next, the mixture was sonicated (20% power, 6 cycles by 4-sec pulse / 2-sec pause), cooled down on ice for 2 min, and then sonicated again, using Fisher Sonic Dismembrator (Fisher Scientific). The CysC-loaded BDEVs (cEVs) were then washed to remove free CysC by ultracentrifugation at 100,000×*g* for 70 min. The EV pellet was resuspended in PBS and diluted to 4.8 × 10^9^ particles/µL. As a control (empty EVs; eEVs), BDEVs were not mixed with CysC, but underwent the same procedure as the BDEVs mixed with CysC.

### Oxygen-glucose deprivation (OGD) and CysC (free or loaded in BDEVs) treatment

To mimic the conditions in the penumbra region, where synaptic disruption has taken place while neuronal death has not yet started, OGD was performed on primary neurons for 20 min on day *in vitro* (DIV) 14 as previously described with some modifications [24]. In brief, cells were washed twice with 1 mL of pre-warmed OGD medium (Neurobasal A medium without glucose (Gibco #A24775-01) containing 0.1% Gentamicin (Thermo Fisher #15750-060), 1× Glutamax (Thermo Fisher #35050-038), and 2% B27 serum (Gibco #17504-044)) to deplete glucose from intracellular storage, and then incubated in the pre-warmed OGD medium (1 mL/well). Afterwards, the 24-well plate was put without lid yet under sterile conditions into an anaerobic chamber (Modulator incubator chamber; Billups-Rothenberg), which was flushed with a gas mixture of 95% N_2_ and 5% CO_2_ for 5 min at a flow rate of 4-6 L/min to yield an N_2_-enriched atmosphere before the complete seal. The anaerobic chamber was placed back into the incubator at 37°C for 20 min [25]. After OGD exposure, the cell plate was removed from the anaerobic chamber, and the OGD media was changed back to the normal maintenance medium under sterile conditions. The control group treated in parallel underwent the same procedure (except for the washing step with OGD medium), was maintained in normal culture medium, and incubated under normoxic conditions with 5% CO_2_ at 37°C. Free CysC (human urine CysC; Millipore Sigma #240896-50UG) was dissolved in 100mM sodium acetate buffer pH = 4.5 and added at a concentration of 0.15µM into the maintenance medium after OGD exposure. As a control, the same volume of solvent but without CysC was used. For the experiments involving CysC loaded into BDEVs (cEVs) or empty BDEVs (eEVs, used as a control), we used the amount of BDEVs that contained between 18 ng or 36 ng of CysC per well and added them to the cells after OGD. Thus, four conditions for every set of experiments were set up: normoxia + solvent/eEVs; normoxia + CysC/cEVs; OGD + solvent/eEVs; and OGD + CysC/cEVs.

### LDH and MTT assay

The CyQUANT™ LDH Cytotoxicity Assay (Thermo Fisher #C20300) was used to detect neuronal cytotoxicity 24 h after OGD according to the manufacturer’s manual. As a positive control (100% of cell death), the provided lysis buffer was diluted to 1× with cell medium in the designated wells. The CellTiter 96® Non-Radioactive Cell Proliferation Assay (MTT) (Promega #G4000) was used to check for cell viability 24 h after OGD treatment in the 24-well plate following the manufacturer’s instructions. Wells filled with medium containing only solvent were used as background control. Absorbance was recorded at 490 nm and 570 nm for the LDH and MTT assay, respectively, using a BioTek microplate reader. Two to three replicates were made for each group.

### Immunocytochemistry and confocal microscopy

Immunocytochemistry was performed as previously described [26]. Briefly, coverslips were fixed with 4% PFA at RT for 10 min and then permeabilized with 0.5% saponin in PBS for 10 min. Next, the coverslips were incubated with 1% BSA in PBS for 30 min at RT to block unspecific binding, further incubated with primary antibodies overnight, and, after 3 washes in TBST, lastly incubated with the corresponding secondary antibodies in 0.1% BSA in PBS. The primary antibodies used here were: goat against MAP2 (S-15; 1:100; Santa Cruz #sc-12012) and rabbit against Synapsin1 (1:500; Synaptic Systems #163103). Secondary antibodies were: anti-goat 555 (1:500; Thermo Fisher #A21432) and anti-rabbit 488 (1:500; Thermo Fisher #AF21206). Images were taken with a Leica TSC SP8 confocal microscope to quantify synaptic density. Roughly 10 neurons (ranging between 7-13) were randomly chosen per coverslip, and 2-3 coverslips were checked per condition per experiment, thus a total of 20- 25 neurons per condition per experiment were imaged. Pictures were taken with the 63× objective with 1× digital zoom. For the 3D images, z-stacks were set up at 0.3 µm. The starting and ending points of the z-series were determined by dendritic spatial distribution on the z-axis. Only neurons that were clearly isolated but also not far away from other neurons and with a dendritic length between 80-100% width or height of the picture frame (90 x 90 µm) were chosen. For synaptic quantification, the z-stacks were reconstructed with Imaris software (Bitplane 7.4.2). The optimized settings used to reconstruct the dendrites and synapses within the surpass module were fixed for every experiment. Synaptic density was considered as the number of Synapsin1 positive puncta per unit area of the dendrites, indicated by the MAP2 fluorescence signal revealing neuronal shape.

### Intracerebroventricular (ICV) injection of BDEVs

Mice were subjected to tMCAO as described above, and 6 h after artery occlusion and reperfusion, 2 µL of resuspended BDEVs loaded or not with CysC were injected into the ipsilateral ventricle with a stereotactic device (Stoelting #51730D). The injection coordinates used were 1.1 mm lateral, 0.5 mm posterior, and 2.3 mm ventral to the skull Bregma. All the experiments were performed blindly as the tubes were labelled with A or B and the experimenters did not know which ones contained BDEVs loaded with CysC or empty BDEVs.

Mice were sacrificed 24 hours after ICV injection with BDEVs, their brains were removed, and the infarct area along with the surrounding penumbra were excised and homogenized in HEPES buffer (10mM, pH = 7.4, supplemented with P.I.) on ice. The homogenates were kept at −80°C. After measuring total protein concentration by BCA assay, samples were used for proteomics or western blot analysis.

### TTC cell viability assay staining

2,3,5-Triphenyltetrazoliumchlorid (TTC) solution was used to confirm the overall stroke volume. The colorless TTC is converted by the mitochondrial succinate dehydrogenase of living cells into formazan, turning viable brain tissue into red, while the dead tissue areas (i.e., the stroke core) remains white. Mouse brains were sliced rostrocaudal into serial 1 mm thick slices and the whole set of brain slices was incubated in the TTC solution for 20 min at RT. Brain slices were scanned and the stroke volume was calculated with Image J software (National Institutes of Health, NIH).

### Immunohistochemistry and microscopy analysis

Brain tissues were fixed with 4% PFA at 4°C for 24 h, rinsed in ice-cold PBS, and embedded in paraffin in a Modular Tissue Embedding Center (Leica #EG1150H). Paraffin-embedded sections were cut in 5 µm thick slices with a microtome (SM 2010R Leica) and affixed to Superfrost microscope slides (VWR #631-0108). After drying at 37°C overnight, slides were deparaffinized and boiled for 20 min in citrate buffer (0.1M sodium citrate and 0.1M citric acid in distilled H_2_O) for antigen retrieval and incubated with 5% normal rabbit serum (Vector laboratory #S-5000) in PBS containing 0.2% Triton for 1 h at RT to block unspecific binding. Afterwards, the sections were incubated with primary antibodies diluted in 5% normal rabbit serum in PBS at 4°C overnight in a wet chamber. The next day, slides were recovered to RT for 30 min and, after four PBS washes of 5 min each, incubated with secondary antibodies diluted in 5% normal rabbit serum in PBS for 45 min at RT in the dark. Samples were finally mounted with DAPI Fluoromount-G mounting medium (Southern Biotech #0100-20). Primary antibodies used were: NeuN (1:2,500; EMD Millipore #ABN91), GFAP (1:100; Invitrogen #13-0300). Secondary antibody used included: anti-chicken AF647 (1:500; Thermo Fisher #A21449), anti-rat AF555 (1:500; Abcam #ab150154). Anti-Isolectin GS-IB4, Alexa Fluor 488 (Invitrogen #121411) antibody was applied with the secondary antibodies. Immunofluorescence imaging was conducted with an Apotome microscope (Zeiss) with a 20× objective.

### Data processing and statistical analysis

For the analysis of LC-MS-derived Proteome data, each experiment was handled separately. Protein abundances were loaded into Perseus (Max Plank Institute for Biochemistry, Version 2.0.3) and log2 was transformed to approximate the Gaussian probability distribution. *T*-testing was performed, accepting proteins identified at a *p*-value < 0.05 with a fold change difference of > 1.5 being considered as significantly differentially abundant between sham and stroke mice. *T*-testing results were visualized using an in house-script, implementing the ggplot2 and ggrepel packages in the R-software environment (version 4.4.2). For functional analysis, proteins found at least 3 times in at least one group (sham and stroke) for at least one time point were considered. To obtain complete data matrices, random forest imputation was performed based on log2 transformed, centered data, using the Random Forest package, integrated into the R software environment (version 4.4.2.; 1,000 trees, 10 repetitions). REACTOME-based Gene Set Enrichment Analysis (GSEA) was performed by using the GSEA software (version 4.3.2; Broad Institute, San Diego, CA, USA), and 1,000 permutations were used. Permutation was performed based on gene sets. A weighted enrichment statistic was applied using the signal-to-noise ratio as a metric for ranking genes. Meandiv scaling was applied. As in default mode, gene sets smaller than 15 and bigger than 500 genes were excluded from the analysis. For visualization of GSEA results, the Enrichment Map (version 3.3) application within the Cytoscape environment (version 3.8.2) was used. Gene sets were considered if they were identified at an FDR < 0.25 and a *p*-value < 0.1. For gene-set-similarity filtering, data set edges were set automatically. A combined Jaccard and Overlap metric was used, applying a cut-off of 0.375. For gene set clustering, Auto Annotate (version 1.3) was applied, using the Markov cluster algorithm (MCL). The gene-set-similarity coefficient was utilized for edge weighting.

For Principal Component Analysis (PCA) we performed linear PCA based on all proteins found in all samples. Log2 transformed, and normalized values were used. To cope with missing values, imputation under the missing not at random (MNAR) hypothesis was performed, using the mean value per column, with a downshift of 1.8. To avoid data distortion in statistical testing, imputation was performed after *T*-testing, for significant candidates only. The first two main principal components were plotted against each other, using a scatter plot in Perseus (version 2.0.3). Hierarchical Clustering was performed, using the Pearson correlation as a distance metric for clustering. For linkage, the average linkage was used. Before clustering, log2 transformed, normalized protein abundances were centered around the mean value per protein for visual purposes.

GraphPad Prism 9 software was used to analyze the data from western blots, cellular viability and cytotoxicity, immunocytochemistry, and immunofluorescence experiments. Student’s *t*-test or Mann-Whitney U test was used to compare the difference between groups, and the statistical difference was considered when \**p* < 0.05, \*\**p* < 0.01, \*\*\**p* < 0.001. Values are described as mean ± SEM. The exact *p* values are given in the text.

## RESULTS

### Synaptosome characterization

Our first goal was to analyse protein changes at the level of the synapses in the core and penumbra area at different time points after tMCAO in mice. For this, we excised the affected area of the ipsilateral hemisphere and isolated the synaptosomes as shown in Fig. 1A. Western blot analysis of the synaptosomal fraction (SN) revealed enrichment of several synaptic proteins (Synapsin1, PSD95, and SNAP25) when compared to a total homogenate of the brain (TH) (Fig. 1B). The SN fraction lacks the nucleic marker Histone H1 and the Golgi marker GM130, used here as indicators of potential contamination from other cellular compartments (i.e., negative control). A representative electron microscopy picture in Fig. 1C shows several structures attributable to bona fide synaptosomes (highlighted in the squared boxes).

### Cystatin C is upregulated in the acute phase of stroke

Ito *et al.* [14] described that, after tMCAO was performed in Mongolian gerbils, axons and dendritic spines decreased till day 4 to eventually start recovering from this time point onwards. To investigate the temporal proteomic changes of the neuronal synapses after stroke in a mouse model, we isolated the synaptosomes from tMCAO- and sham-treated mice at 24 hours (shams: n = 4; tMCAO: n = 3), 4 days (shams: n = 5; tMCAO: n = 4) and 7 days (n = 5 per group) after stroke and performed mass spectrometry-based proteome analysis. At 24 h we identified 144 proteins significantly differentially abundant in stroke compared to sham with a *p*-value ≤ 0.05. Out of these, 85 (30 upregulated, 55 downregulated in stroke) exceeded the log2 fold-change (log2FC) cutoff of 1.5. After 4 days, 195 proteins were significantly differentially abundant between shams and strokes (*p*-value ≤ 0.05). Out of these, 81 exceeded the log2FC ≥ 1.5 (43 upregulated in stroke, and 38 downregulated). Finally, at 7 days post insult, 173 proteins were found significantly differentially abundant in stroke compared to shams (*p*-value ≤ 0.05) with 23 proteins being significantly upregulated and 43 significantly downregulated log2FC ≥ 1.5. Heat maps and supervised principal component analysis (PCA) showing the differential clustering of the sham samples and the tMCAO samples are shown in Suppl. Fig. 1. The volcano plots in Fig. 2A, 2C, and 2E present the names of the top up-and downregulated proteins for each time point, whereas Fig. 2B, 2D, and 2F depict the changes between sham and strokes at different time points analysed by REACTOME-pathway-based GSEA. This analysis showed that changes at 24 h comprised upregulation of biological pathways related to Cell Cycle, Translation, and Golgi-Associated Trafficking in the tMCAO group, whereas biological pathways related to NMDA Receptor Signalling or Ion Channel Homeostasis were decreased in tMCAO compared to shams. At 4 days after stroke, pathways associated with NMDA Receptor Signalling or Ion Channel Homeostasis were still decreased in the tMCAO group compared to shams whereas proteins associated with Cell Cycle, Translation, Golgi-Associated Trafficking, Autophagy or Lipid Metabolism pathways showed significant upregulation in the tMCAO group. Finally, at 7 days, no concrete pathway could be associated to be overrepresented in the tMCAO group, whereas pathways related to energetic metabolism such as Carbohydrate Metabolism or Citric Acid Cycle pathways showed significant overrepresentation in the sham group (thus, were decreased in tMCAO mice).

**Figure 2.**
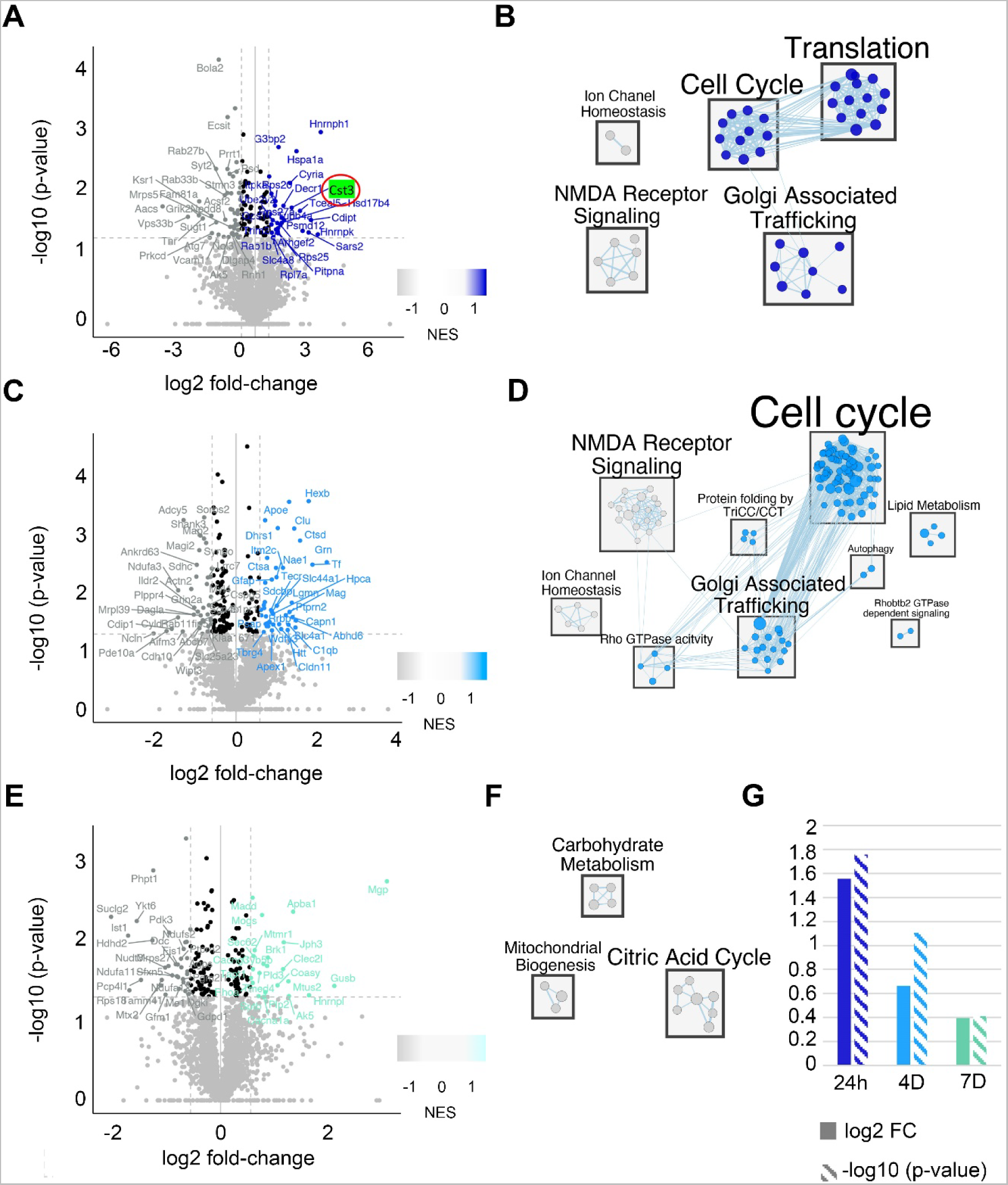
Proteomic analysis reveals upregulation of CysC at 24 h after stroke in synaptosomes. (A) Volcano plot showing in dark blue the significantly upregulated proteins in synaptosomes 24 h after stroke and in grey the significantly downregulated proteins. Highlighted is Cst3 (CysC). (B) GSEA based on the proteins found to be significantly regulated a 24 h after stroke. Translation, Cell Cycle, and Golgi-associated Trafficking (dark blue dots) are the most enriched processes in stroke, indicating that, at this time point, synaptic reorganization is taking place. NMDA Receptor Signaling and Ion Channel Homeostasis (light grey points) are pathways overrepresented in shams, therefore decreased in the stroke group. (C) Volcano plot showing in light blue the significantly upregulated proteins in synaptosomes 4 d after stroke and in grey the significantly downregulated proteins. (D) GSEA based on the proteins found to be significantly differentially regulated at 4 d in synaptosomes after stroke. Again, Cell Cycle, NMDA Receptor Signaling, and Golgi-associated Trafficking are the more enriched sets. However, at this time point, some other sets such as Lipid Metabolism of Autophagy are also enriched in the tMCAO group (light blue dots) compared to shams. (E) Volcano plot showing in light green the significantly upregulated proteins in synaptosomes 7 d after stroke and in grey the significantly downregulated proteins. (F) GSEA for the 7 d time point. Enrichment of protein sets were only found in shams (grey dots) and related to metabolic processes (Citric Acid Cycle, Carbohydrate Metabolism) and Mitochondrial Biogenesis. (G) Graph showing the log2FC and the −log10 (*p*-value) for CysC at the different time points of the study. At 24 h, the log2FC is 1.55 and the *p*-value is 0.017. At 4 d the log2FC is 0.66 and the *p*-value is 0.07 and, thus, slightly above the *p*-value cutoff (*p* < 0.05). After 7 d, no significant differences could be detected between shams and strokes regarding CysC (Cst3) (log2FC = 0.4; *p* = 0.39).

For the present study, our working hypothesis was that proteins known to be protective or beneficial in ischemic injury and found upregulated at early time points after stroke could represent candidates involved in an endogenous mechanism of response to ischemic damage, worth to be studied and employed as a potential therapeutic option. Among the upregulated proteins at 24 h after tMCAO, we found that cystatin C (CysC, CST3), a protease inhibitor known to exert protective effects against ischemic brain injury and neurodegeneration [27, 28] was significantly (*p* = 0.032) upregulated by 2.9- fold in the stroke synaptosome preparations. As depicted in Fig. 2G, CysC was still increased at 4 d after ischemia/reperfusion (I/R) compared with the sham group by 1.5-fold but showed a higher heterogeneity in stroke animals, most likely related to differences in the speed of neuronal recovery between individual animals (*p* = 0.07). After 7 days, CysC levels in the I/R group were found to be comparable to the sham group.

To confirm the results obtained for CysC by quantitative mass spectrometry, we performed western blot analysis. As shown in Fig. 3, expression levels of CysC in the tMCAO group were significantly elevated (*p* = 0.0286) at 24 h post-stroke and, with this technique, also at 4 days after stroke (*p* = 0.0159), while this difference between groups disappeared 7 days post-stroke. Notably, expression levels of the synaptic markers SNAP25 and PSD95 at 24 h post-stroke were significantly lower in the tMCAO group, possibly indicating a substantial loss of synapses at this time point (both *p* = 0.0286). This particular difference was not seen later, at 4 and 7 days post-stroke, maybe indicating a partial synaptic recovery over time. Thus, fitting in our premises for a candidate protein, we decided to focus on CysC to further study its effects on synapses under ischemic damage.

**Figure 3.**
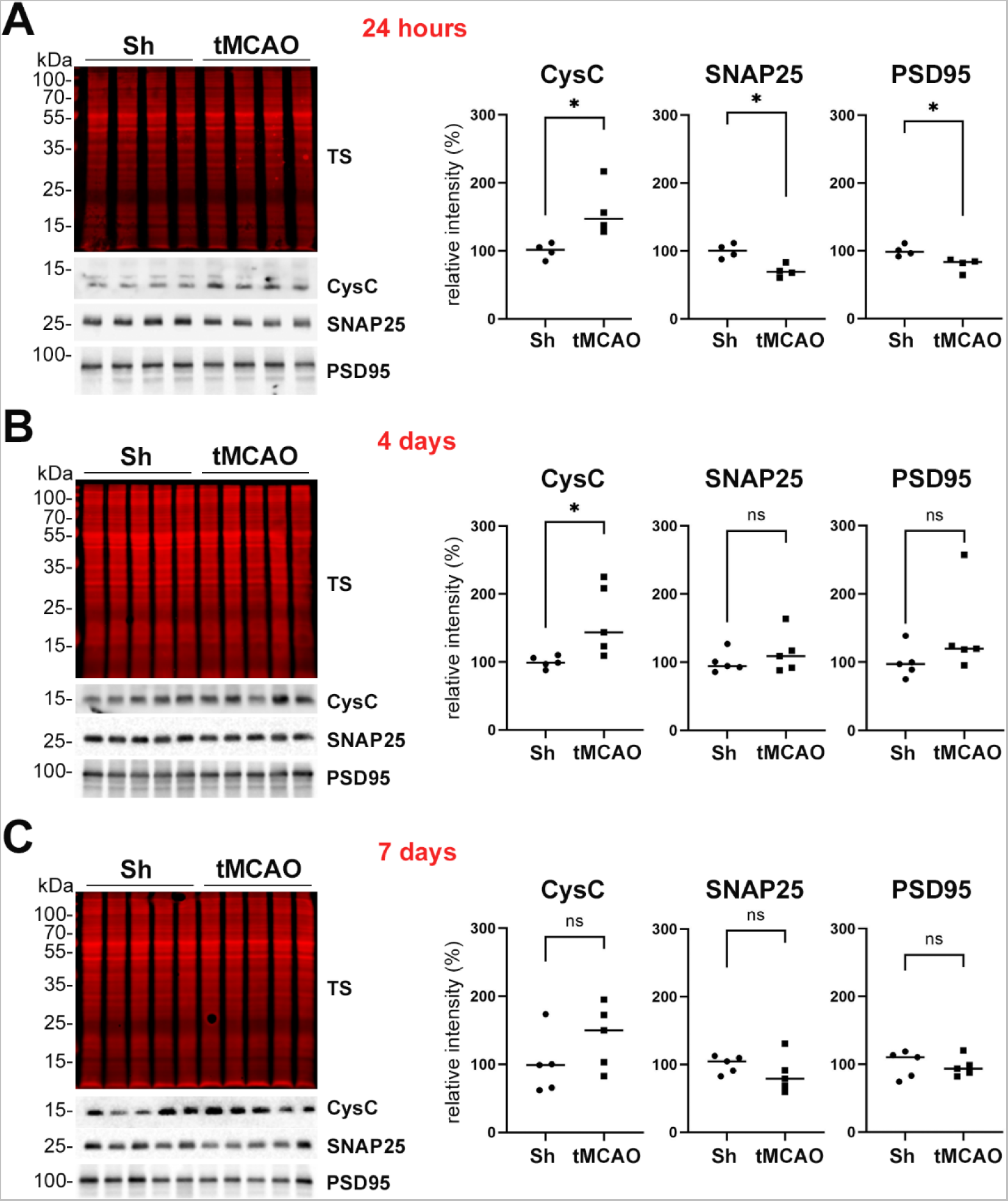
CysC is enriched in synaptosomes at 24 h and 4 d after stroke when assessed by western blot. (A) Western blot of synaptosome-enriched fractions of shams (Sh) and stroked mice (tMCAO) for CysC, SNAP25, and PSD95. TS is a total protein staining showing equal loading for each sample. On the right, dot plot graphs of the relative intensity quantification of the western blot bands reveal that CysC expression is upregulated in the tMCAO group at 24 h, whereas SNAP25 and PSD95 are significantly downregulated at this time point after tMCAO. (B) Same analysis as in (A), but at 4 d after tMCAO. Dot plot graphs on the right show that CysC is still significantly upregulated, whereas SNAP25 and PSD95 show similar levels when compared to shams. (C) Same analysis as in (A) and (B), but for 7 d after tMCAO. Dot plot graphs show that there are no significant differences among any of the proteins analyzed between shams and strokes at this time point. \**p* < 0.05; ns: not significant.

### CysC, either free or loaded into BDEVs, has a protective effect on synapses of neurons subjected to OGD

To test the effect of CysC on hypoxia-challenged neurons *in vitro*, we first established the proper conditions simulating ischemic damage in the penumbra area, where neurons are still viable and can be rescued. For this, based on our previous experiences in the lab, we submitted primary neurons isolated from P0-P1 mice to oxygen-glucose deprivation (OGD) treatment for 20 min. To additionally measure whether the addition of CysC was causing any effect on neuronal viability, we established four experimental groups (normoxia or OGD, and each either with or without CysC administration), and cytotoxicity and viability were assessed with LDH and MTT assays, respectively. As shown in Fig. 4A, no significant differences in cytotoxicity or viability were observed between the OGD and normoxia groups, implying that the 20 min OGD treatment only caused negligible cell death. The addition of CysC did not have any effect either in normoxia nor after these OGD conditions.

**Figure 4.**
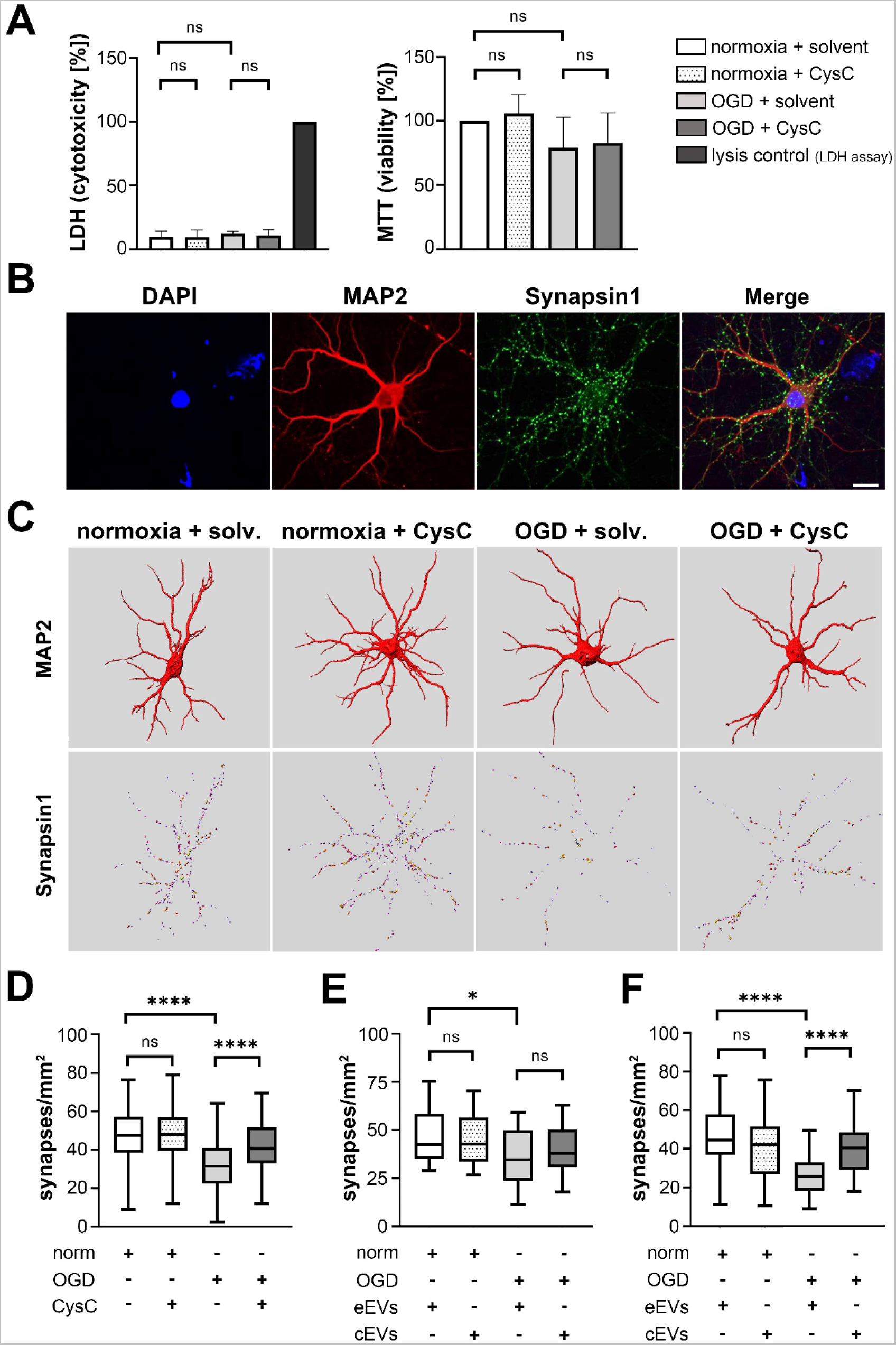
CysC, either free or encapsulated in BDEVs, rescues synapses in neuronal cultures after ischemic insult *in vitro*. (A) LDH toxicity assay on the left and MTT viability assay on the right show that, after 20 min of OGD/reperfusion, there are no differences in the survival of primary neurons treated or not with CysC compared to normoxic conditions; ns: no significance. (B) Representative confocal microscopy pictures (63× objective) showing dendritic staining with MAP2 in red and synaptic terminals labelled with Synapsin 1 in green. This staining was used for the quantification of the synapses. DAPI in blue is used as a nuclei marker. Magnification is 1×, scale bar is 10 µm. (C) Representative three-dimensional reconstruction of the dendritic branches and synaptic puncta of individual neurons by Imaris software used to quantify the synapses. It can already be observed that, after 20 min of OGD, many synaptic structures disappear but are preserved after treatment with free CysC. No effects of free CysC were observed under normoxic conditions. Solv: solvent. (D) Graph showing the quantification of synapses/mm^2^ under normoxic conditions and OGD conditions with and without free CysC. While the addition of CysC has no effect under normoxic conditions, the addition of free CysC significantly increases synaptic puncta. (E) Same analysis as in (D) but with neurons under normoxic or OGD conditions having received treatment with 14 µL of empty BDEVs (eEVs) or 14 µL of CysC-loaded EVs (cEVs). The quantification shows no significant differences between the treatments neither in normoxic nor under OGD conditions. (F) Same analysis as in (E) but after delivery of the double amount of eEVs or cEVs (28 µL). Neurons subjected to OGD and treated with 28 µL of cEVs significantly improved the number of synaptic structures compared to the non-treated ones, showing a dose-dependency of cEVs treatment. Norm: normoxia; eEVs: empty EVs; cEVs: CysC-loaded EVs; \**p* < 0.05; \*\**p* < 0.01; \*\*\**p* < 0.001; \*\*\*\**p* < 0.0001; ns: not significant.

We then proceeded to check the status of the synapses in these four conditions. For this, we labeled the neurons with MAP2 (labelling the dendrites of mature neurons) and Synapsin1 (a resident protein of synaptic vesicles) as a proxy for axon numbers (Fig. 4B and Suppl. Fig. 2). After OGD or normoxic conditions, neurons were treated with or without CysC and stained. More than 100 neurons for each condition (normoxia + solvent (n = 114), normoxia + CysC (n = 113), OGD + solvent (n = 114), OGD + CysC (n = 114)) were analysed with the Imaris program (Fig. 4C). As shown in Fig. 4D, neuronal synaptic density was significantly reduced after OGD treatment (OGD + solvent: 31.88 synapses/1000 µm^2^ ± 12.56; *p* < 0.0001) when compared to normoxia conditions (normoxia + solvent: 46.96 synapses/1000 µm^2^ ± 12.55), while the addition of CysC after OGD significantly increased synaptic density (OGD + CysC: 42.00 synapses/1000 µm^2^ ± 11.58; *p* < 0.0001). The addition of CysC did not change the synaptic density of neurons in the normoxia group (normoxia + solvent: 48.60 synapses/1000 µm^2^ ± 11.96, *p* = 0.3155). It was recently described that CysC loaded in EVs conferred protection against nutrient deprivation in primary neurons [29]. EVs have the potential to be used as therapeutic vehicles for drug delivery and treatment of neurological diseases, as they (i) efficiently cross the blood-brain barrier (BBB) [30], (ii) increase drug biostability as they protect their cargos from extracellular degradation crediting to their specific membrane composition [19], and (iii) have high biocompatibility and low immunogenicity [31, 32]. Thus, we checked whether the effect seen with free CysC could be reproduced when CysC was loaded into BDEVs as a proof of concept for future treatment approaches. After OGD or normoxic conditions, primary neurons were treated with either non-loaded (described here as “empty” BDEVs, eEVs) or BDEVs loaded with CysC (cEVs), and the same analysis was performed as described above. Two doses of cEVs were studied: 14 µL (4.8 × 10^9^ particles/µL, 1.28 ng/µL of CysC, total amount of 17.92 ng of CysC; n = 20 neurons per condition) or 28 µL of EVs (equivalent to 4.8 × 10^9^ particles/µL, 1.28 ng/µL of CysC, total amount of 35.84 ng of CysC; n = 60 neurons per condition)

While no difference in treatment of cEVs in normoxia or OGD was observed for the treatment with 14 µL of cEVs (Fig. 4E), significant differences were observed after adding the double amount of cEVs after OGD treatment (Fig. 4F; *p* < 0.0001, synaptic density after OGD = 26.34 synapses/1000 µm^2^ ± 9.627 and after OGD with cEVs treatment = 40.03 synapses/1000 µm^2^ ± 11.83), indicating a dose-dependent effect of CysC loaded into BDEVs.

### BDEVs loaded with CysC rescue synapses *in vivo* after induced stroke in mice

To assess whether the protective effect of CysC on synapses observed *in vitro* is also present *in vivo*, we injected eEVs or cEVs intracerebroventricularly (ICV) 6 h after tMCAO and sacrificed the mice 24 h afterwards to evaluate treatment outcome. We chose this time point as we hypothesized that 6 h post-insult might be relevant for treatment in a clinical setting. To analyse changes in the volume of stroke, we used the TTC cell viability assay with n = 6 per group (Fig. 5A). After quantifying the ratio between stroke area related to total brain area, we found no significant differences in the brain infarct area between the two groups of mice (Fig. 5B). Immunohistochemistry for NeuN (as a marker of nuclei and cell bodies of most differentiated neuronal cell types in rodents), isolectin B4 (as a marker of rodent cerebral vasculature and microglia) or GFAP (as a marker of astrocytes) likewise showed no significant differences between the two treated groups (Suppl. Fig. 3; eEVs n=6, cEVs n=5).

**Figure 5.**
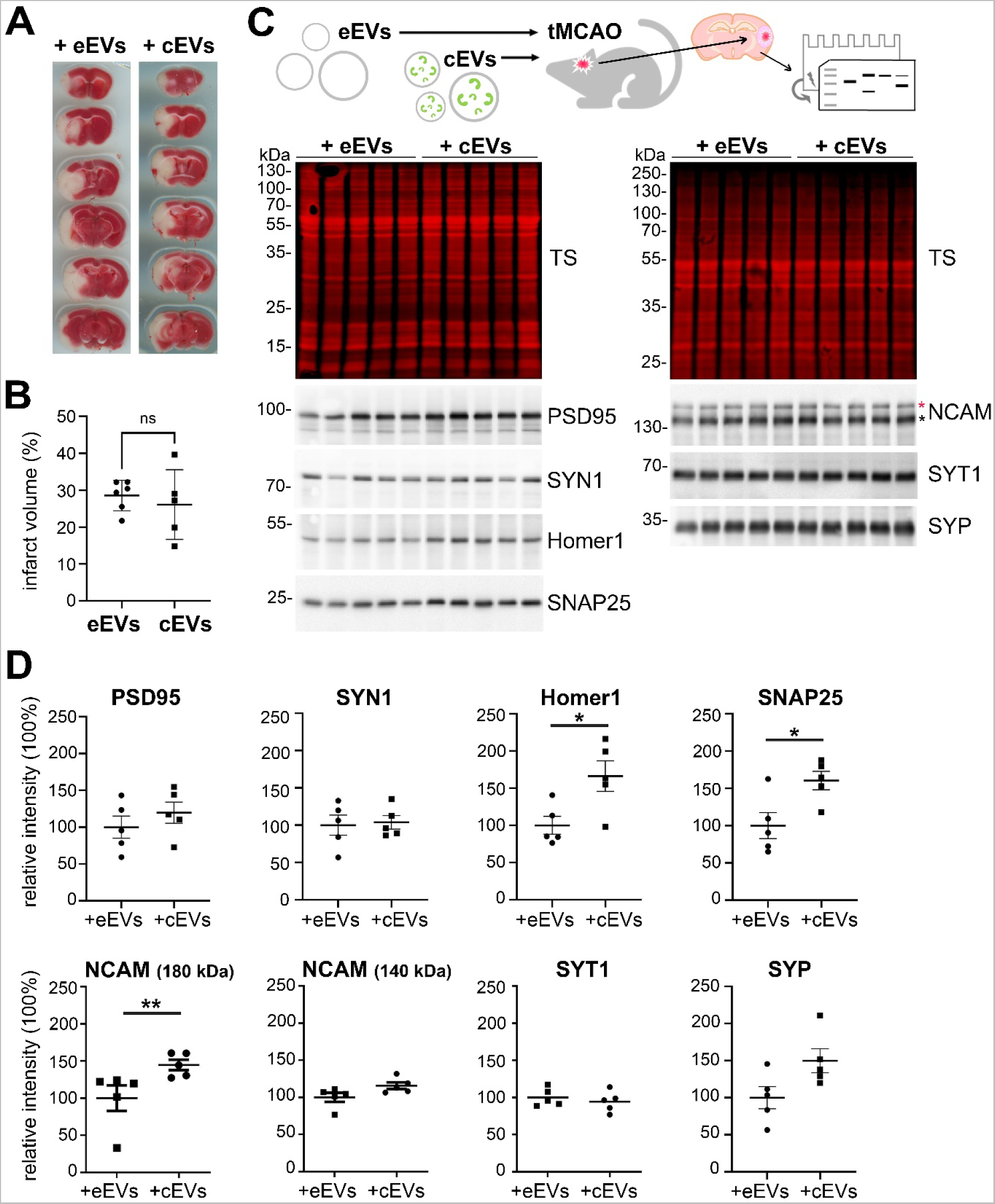
BDEVs loaded with CysC significantly enhance the expression of several synaptic markers 24 h after treatment following tMCAO in mice. (A) Representative images of the TTC staining 30 h after tMCAO (and 24 h after BDEVs delivery) from brains of mice either treated with empty BDEVs (eEVs) or with BDEVs loaded with CysC (cEVs). (B) Dot plot showing the quantifications of the infarct volume ratio of animals treated with eEVs or with cEVs after tMCAO. Although there is a tendency towards a decreased infarct volume in the brains of animals treated with cEVs, this did not reach statistical significance. (C) Brains of animals subjected to tMCAO and then treated either with eEVs or cEVs were analysed by proteomics (Suppl. Fig. 4) and western blot (upper scheme) using different pre- and post-synaptic markers as indicated in the lower panel. TS is total protein staining confirming that similar amounts of sample were loaded. (D) Dot plots showing the quantification of the relative band intensity for each of the synaptic proteins analyzed comparing both groups of animals (tMCAO treated with either eEVs or cEVs). Each band was referred to the same sample lane on the total staining for normalization. PSD95, Synaptotagmin (SYP), Homer1, SNAP25, and NCAM (180 kDa) showed increased amounts in cEVs-treated mice although only the last three reached significance. Synapsin1 (SYN1), NCAM (140 kDa), and Synaptophysin (SYT1) did not show any differences in amounts between animals treated with eEVs or cEVS. eEVs: empty BDEVs; cEVs: CysC-loaded BDEVs; \**p* < 0.05; \*\**p* < 0.01; \*\*\**p* < 0.001; \*\*\*\**p* < 0.0001; ns: not significant.

To explore the pathways activated by the treatment with cEVs, we performed mass spectrometry of the ipsilateral hemisphere of the eEVs- (n = 5) and the cEVs-treated groups (n = 5). As shown in Suppl. Fig. 3, we identified 3,913 proteins, 6 of which were significantly upregulated (log2FC ≥ 1.5; *p* ≤ 0.05) in cEVs, and 9 significantly downregulated (log2FC ≤ −0.58; *p* ≤ 0.05). No enrichment cluster analysis was possible due to the low amount of upregulated proteins, however, five out of the six upregulated proteins (Developmentally-regulated GTP-binding protein 2 (DRG2); Exopolyphosphatase PRUNE1 (PRUNE1); Craniofacial development protein 1 (CFDP1); Cullin-1 (CUL1); and Centrosomal protein of 170 kDa protein B (CEP170B)) can be related to either microtubule reorganization or cell proliferation. Finally, because it was very difficult to assess and reliably quantify synaptic markers by immunohistochemical analysis in brain samples, we performed western blotting (n = 5 for each group) for several known pre- and post-synaptic markers. Homer-1 and PSD95 are components of the postsynaptic density, while SNAP25, Synaptophysin, Synaptotagmin, and Synapsin1 are used as presynaptic markers. NCAM, a neural cell adhesion molecule implicated in synaptic plasticity, has three isoforms with two of them (NCAM 180 kDa and NCAM 140 kDa) being located at the pre-and post- synaptic membrane, while the third isoform (NCAM 120 kDa) is mostly expressed in glia [33]; we therefore quantified the 180 and 140 kDa bands relevant for neuronal synapses. As shown in Fig. 5C and 5D, while expression levels of Synapsin1 and Synaptotagmin did not show any upregulation, Homer-1, SNAP25, and NCAM 180 kDa were significantly upregulated in the cEV-treated group compared to the control group (*p* = 0.031, *p* = 0.031 and *p* =0.008, respectively). Levels of Synaptophysin, PSD95, and NCAM 140 kDa showed an increased trend in the cEV treatment group, yet this did not reach statistical significance (*p* = 0.0952, *p* = 0.4206 and *p* = 0.0556, respectively). Thus, we conclude that also *in vivo*, CysC-loaded BDEVs confer beneficial, synapse-protecting or -preserving effects after stroke.

## DISCUSSION

In the present study, as depicted in the summary in Fig. 6, we show that several proteins are upregulated in surviving synapses in the acute phase of stroke, possibly indicating an attempted protective response against the ischemic damage. Among these increased proteins, we chose to study the role of CysC, as it was transiently upregulated at 24 h and 4 days after reperfusion to finally decrease to normal levels after 7 days. When CysC was externally delivered, either free or encapsulated in EVs, it had positive effects at the synaptic level *in vitro* and *in vivo*. As synaptic preservation is fundamental for neuronal survival and functional outcome, treatment with CysC encapsulated in EVs may open the possibility of its use as a targetable therapeutic agent with improved pharmacodynamic and kinetic features.

**Figure 6.**
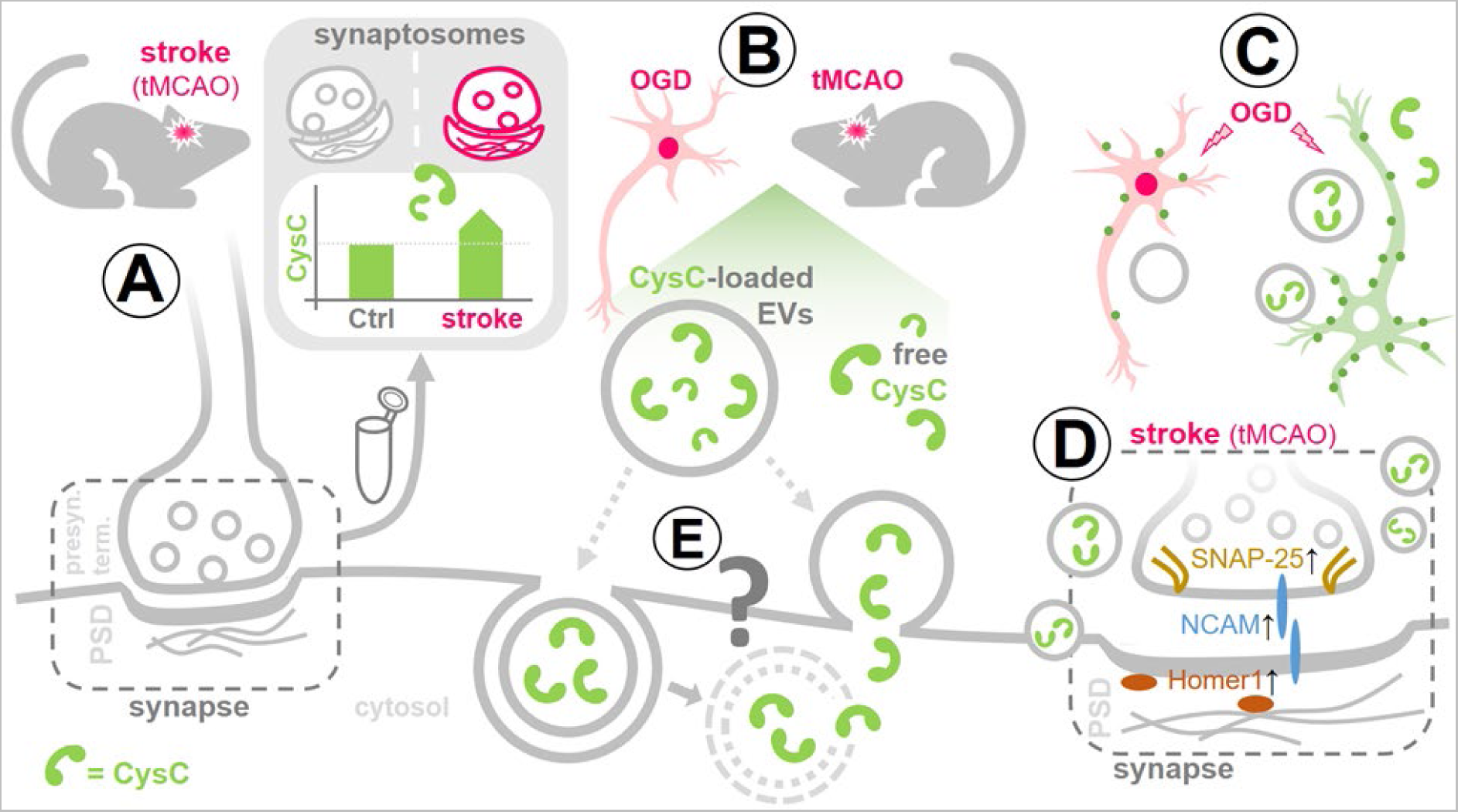
Summarizing scheme. (A) To mimic stroke pathophysiology, mice were subjected to transient middle cerebral artery occlusion (tMCAO). Synaptosomes, i.e. structures consisting of a presynaptic terminal and the attached postsynaptic density (PSD), were isolated from the brains of stroked mice and controls at different time points after the insult. Mass spectrometry analysis of these samples revealed that the cysteine protease inhibitor Cystatin C (CysC) is significantly upregulated in the acute phase after stroke. (B) CysC, either free or loaded into brain-derived extracellular vesicles (BDEVs), was administered to primary neurons after oxygen-glucose deprivation (OGD); and BDEVs loaded or not with CysC were injected into stroked mice (6 h after tMCAO) and controls. (C) OGD-challenged neurons treated with free CysC or CysC-loaded BDEVs (green neuron) had a higher amount of synaptic punctae (green dots) than neurons not treated with any CysC (pink neuron), indicating protective synaptic preservation and/or recovery by CysC. (D) Fittingly, when stroked mice were injected with CysC-loaded BDEVs, pre- and postsynaptic marker proteins were increased in their brains compared to mice that received BDEVs without CysC. (E) The mechanism of cellular uptake and the destiny of CysC-loaded EVs (grey question mark) as well as the mode of action of CysC remain unknown and need to be investigated

Synaptic transmission is the most energy-consuming process of neurons [34] and, thus, after ischemic damage and the subsequent lack of energy, is one of the first processes to be shut down in the penumbra area [17]. In this region, loss of synapses is not always accompanied by neuronal death and, in fact, the rapid distortion suffered by synaptic structures can be fully recovered in case of intervention till 3 h after reperfusion (as observed with two-photon microscopy), suggesting a high capacity for a fast reorganization in this area [16, 35, 36]. Moreover, it was described that, during the recovery phase, the peri-infarcted area shows extraordinary neuronal plasticity, which, in global ischemic models of stroke, could be observed as soon as 4 days after reperfusion [14, 37]. Therefore, it was hypothesized that preserving synaptic structures or promoting their remodelling at the penumbra in a determined time window would improve functional neurological recovery in patients suffering from stroke [38]. Interestingly, in the synaptosomal preparations studied here, gene set enrichment analysis showed an upregulation of cytosolic ribosomal proteins and proteins related to the cell cycle as soon as 24 h after reperfusion, with the latter even being further increased after 4 days. Most of the proteins belonging to the Cell Cycle term were related to ubiquitination and proteasome degradation, which may have dual (i.e., harmful but also protective) roles after ischemic damage, for example by regulating autophagosome activity [39]. Interestingly, some other cell cycle-related proteins, such as Cyclins, can have other functions in neurons, such as regulation of the cytoskeletal organization [40]. This implies that the synaptic recovery starts already at early time points (24 h and 4 d) in our ischemic model.

Among the upregulated proteins found in synaptosomes isolated from stroked mouse brain, we turned our attention to CysC, a secreted cysteine protease inhibitor expressed in the brain by neurons, astrocytes, and microglia [41] and shown to protect neurons from various toxic stimuli, such as nutrient deprivation or oxidative stress *in vitro* [42]. *In vivo*, CysC also confers protective effects after focal ischemia, as CysC knock-out mice presented with enlarged stroke volume. However, in the same study it was also shown that, by using the model of global ischemic damage, neuronal injury was decreased in certain brain populations [43], highlighting rather complex effects of CysC after ischemic insults. By studying preconditioning –the mechanism whereby a mild ischemic insult triggers molecular mediators that protect the brain from future acute ischemic stroke events [44–46]– it was found that, after repeated hyperbaric oxygen (HBO) pre-conditioning in rats followed by tMCAO, one of the molecules significantly upregulated in a serum proteomic screening was CysC [47]. As preconditioning is a paradigm to study lasting responses set up to reduce ischemic damage, the upregulation of CysC would imply an induced endogenous protection response towards stroke. In their model, the authors also observed that CysC was increased at the penumbra as early as 3 h upon insult [48]. In another study, the same group (with the same model) demonstrated that exogenous ICV injection of CysC 30 min after reperfusion leads to a reduction of the infarct volume found at 3 days after stroke [47]. Finally, Yang *et al*. also showed that intraventricular injection of CysC 30 min before tMCAO in mice (mimicking a preconditioning effect) reduced the infarct size 24 h after stroke/reperfusion injury.

Different mechanisms through which CysC exerts its neuroprotection have been proposed [28]. In stroke, it has been described that CysC leads to the preservation of the integrity of the lysosomal membrane and the induction of autolysosome formation, leading to autophagy promotion [47, 48]. But it can also act as a potent inhibitor of cysteine proteases such as cathepsin B and L [49], which play a detrimental role in stroke [50], or by promoting BBB integrity [51]. Our study revealed a significant upregulation of CysC in the synapses at the penumbra during the initial phase of stroke in tMCAO mice. Although we did not inspect the lysosomal system, it is known that, following stroke, neurons increase their autophagic activity, which may result in lysosomal dysfunction and destabilization of the lysosomal membrane, associated with perturbed synaptic ultrastructure and plasticity [52]. Thus, as CysC participates in conserving lysosomal membrane integrity and in stimulating autophagosome formation, we hypothesize that synaptic integrity is a consequence of its role in maintaining optimal lysosomal system performance. However, other experiments are needed to elucidate the possible direct mechanism at a molecular level, and we cannot rule out that other indirect or more complex mechanisms (for instance related to microglia) are implicated in these processes [53].

EVs are nanometer-sized membrane-enclosed particles that can transport cargo between cells and organ systems [54]. EVs were shown to have therapeutic potential [55] in several diseases, including cancer [56], neurological [57], and cardiovascular diseases [58]. The advantages of using EVs as drug vehicles for the central nervous system (CNS) are (i) their size, (ii) their stability in circulation, (iii) their ability to cross biological barriers such as the BBB, (iv) their ability to utilize endogenous cellular uptake mechanisms, and (v) their low immunogenicity [59]. In the present study, we have delivered CysC- loaded BDEVs to stroked mice and demonstrated that at 24 h after ischemia/reperfusion (I/R) some synaptic proteins in both pre- (SNAP25, NCAM) and postsynaptic (Homer1) terminals show increased abundance in treated animals compared to animals that only received empty BDEVs. *In vivo,* synaptic remodelling has been observed in several instances after ischemic damage, but in the acute phase of stroke, most synaptic proteins are decreased. For example, Marti *et al.* showed decreased Synapsin1 immunoreactivity after transient forebrain ischemia in Mongolian gerbils, followed by increased expression peaking at 7 days after stroke [60]. Upregulation of SNAP25 was found to be a synaptic response to ischemic damage in the hippocampus of gerbils two days after stroke [15], while others have found a prominent decrease of SNAP25 and synaptophysin for the same time point in the stroked gerbils that were subsequently incremented at 14 days [61]. In our analysis, we found SNAP25 significantly reduced in isolated synaptosomes at 24 h after stroke. Homer1 is a post-synaptic scaffold protein that promotes dendritic spine stability and has important implications for the whole post-synaptic network, as its absence alters the whole synaptic proteome [62]. It has been reported to be decreased after I/R in rats, probably contributing to the synaptic alterations and the neuronal fate in the penumbra [63]. In our study, treatment with CysC-loaded BDEVs increased these synaptic proteins as soon as 24 h after stroke, suggesting either promotion of synaptic remodelling or synaptic protection by CysC, thus corroborating our *in vitro* results in primary neurons. The fact that, despite protection at the level of synapses, we did not detect a decrease in stroke volume in mice after CysC administration may be attributed to the short time point after reperfusion of our observations (24 h). Longer intervals (e.g., 72 h or 96 h) may shed more light on overall neuronal survival at the penumbra area after CysC treatment, whereas the time point assessed in the present study may just be fitting for detecting changes occurring at the synapse level.

Overall, we show the capacity of CysC to safeguard neuronal synaptic structures against ischemic damage *in vitro* and *in vivo* and the utilization of EVs to effectively deliver CysC. This may open the possibility of its use in therapy as the delivery in EVs probably increases the efficiency of CysC to reach the CNS and prolongs its bioavailability, thus enhancing its protective potential.

## ACKNOWLEDGEMENTS

The authors thank Prof. Dr. Marina Mikhaylova (Humboldt University Berlin) for providing the synaptosome isolation protocol. We also want to thank the UMIF core facility (Dr. Antonio Virgilio Failla) for the support with the confocal microscopy. This study was supported by the Hermann and Lili Schilling Stiftung (to Tim Magnus); the China Scholarship Council (CSC to Yuqi Gui), the Werner Otto Stiftung (to Yuqi Gui); and the National Institute of Health grants AG017617, AG056732, AG057517 and DA044489 to Efrat Levy.

