## Supplemental data for "Cystatin C loaded in brain-derived extracellular vesicles rescues synapses after ischemic insult *in vitro* and *in vivo*"

### SUPPLEMENTAL FIGURES

**Suppl. Fig. 1. Heat maps and principal component analysis (PCA) of the synaptosome proteomic analysis.** Heat maps and scatter plot visualization of the first two principal components in linear PCA, based on all Student's *T*-test significant ($p < 0.05$) proteins between sham and stroke at different time points ((A) 24 h, (B) 4 d, (C) 7 d).

**Suppl. Fig. 2. Representative 3D reconstruction of primary neurons subjected or not to OGD and treated either with eEVs or cEVs.** Dendrites were labeled with MAP2 and synapses with Synapsin 1. The scale bar is 10 µm.

**Suppl. Fig. 3. Microglia and astrocytes show no differences in their immunoreactive pattern between animals treated with eEVs and cEVs.** (A) Representative immunofluorescence pictures of brain slices observed under the Apotome microscope (20×). NeuN (violet) labelled neuronal bodies, whereas the microglia were stained with isolectin B4 (green); astrocytes were labelled with GFAP (red), and nuclei were stained with DAPI. No major differences were observed between both groups. The scale bar is 50 µm.

**Suppl. Fig. 4. Proteomic analysis of brains of mice subjected to tMCAO and treated with either eEVs or cEVs.** (A) Volcano plotting of the -log10 (*p*-value) against the log2FC difference for *t*-testing mass spectrometry results between tMCAO mice treated either with eEVs or with cEVs. (B) Heat map showing differential clustering between tMCAO samples treated with eEVs (red dots) and tMCAO samples treated with cEVs (green dots). (C) Scatter plot visualisation of the two main components in supervised PCA, performed based on all Student's *T*-test significant proteins between tMCAO mouse brains treated either with eEVs or cEVs.

**SUPPL. FIGURE 1**

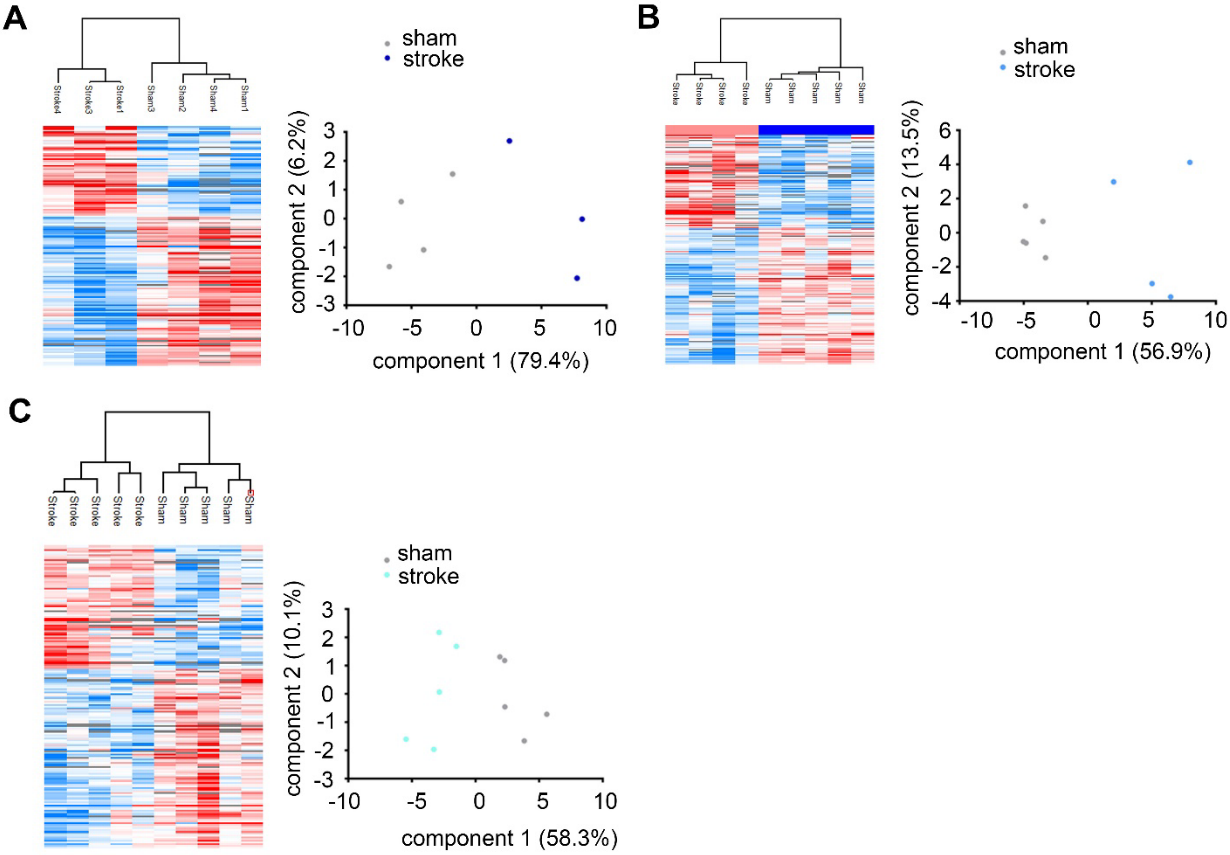


**SUPPL. FIGURE 2**

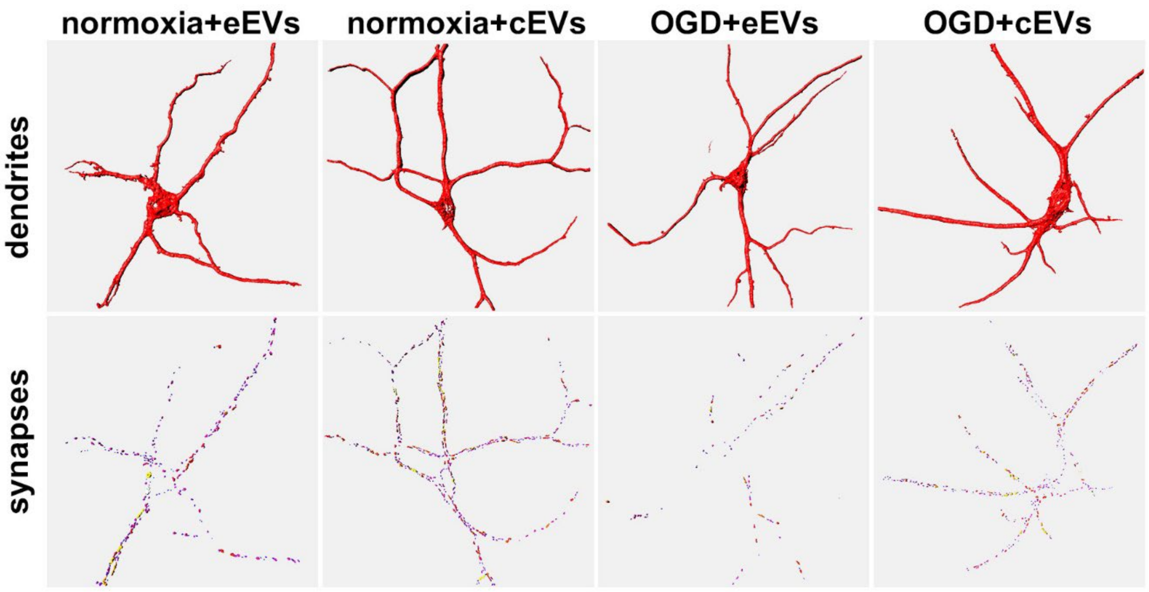

**SUPPL. FIGURE 3**

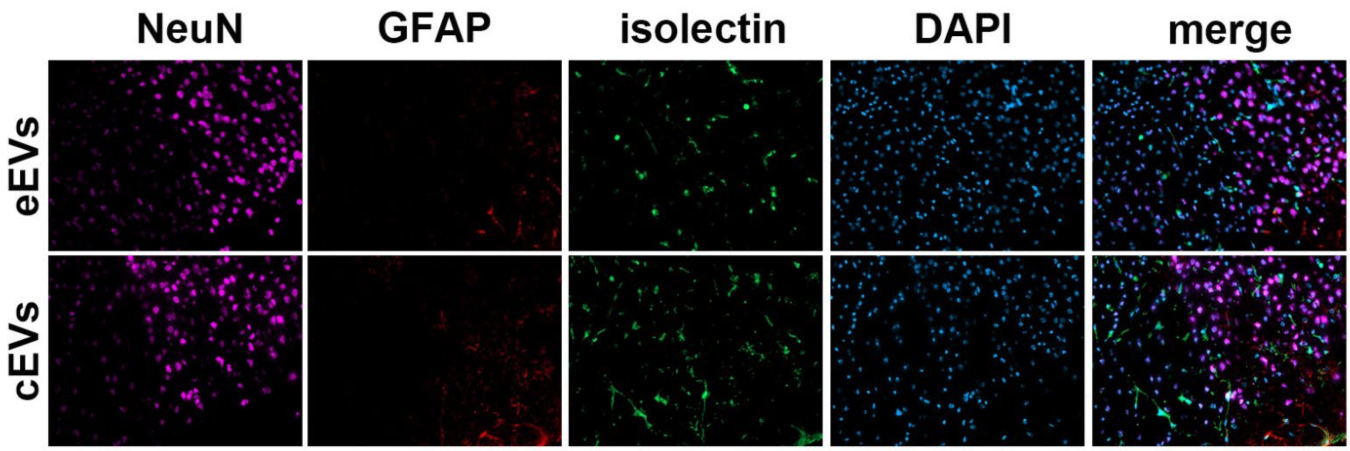


**SUPPL. FIGURE 4**

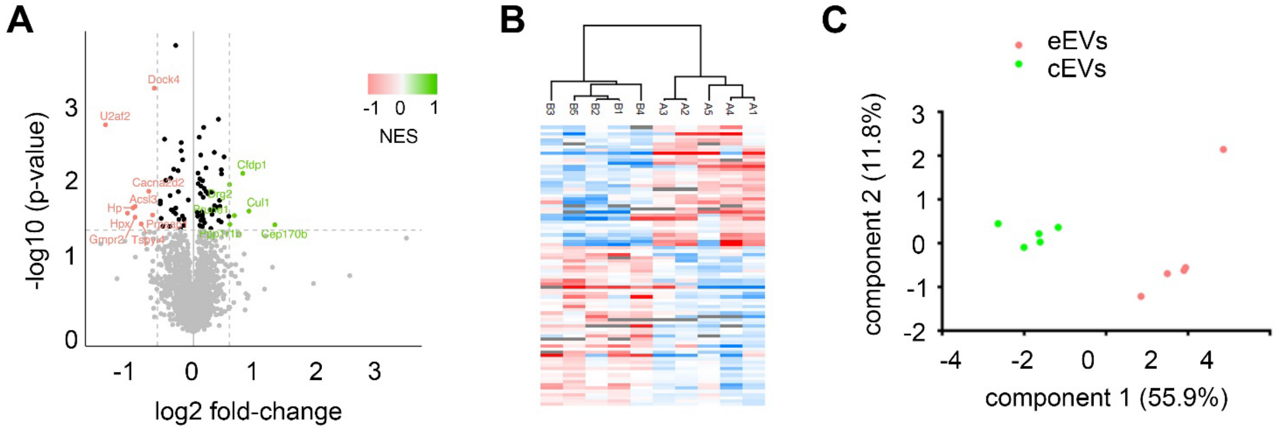
